# Gene flow biases population genetic inference of recombination rate

**DOI:** 10.1101/2021.09.26.461846

**Authors:** K. Samuk, M.A.F. Noor

## Abstract

Accurate estimates of the rate of recombination are key to understanding a host of evolutionary processes as well as the evolution of recombination rate itself. Model-based population genetic methods that infer recombination rates from patterns of linkage disequilibrium (LD) in the genome have become a popular method to estimate rates of recombination. However, these LD-based methods make a variety of simplifying assumptions about the populations of interest that are often not met in natural populations. One such assumption is the absence of gene flow from other populations. Here, we use forward-time population genetic simulations of isolation-with-migration scenarios to explore how gene flow affects the accuracy of LD-based estimators of recombination rate. We find that moderate levels of gene flow can result in either the overestimation or underestimation of recombination rates by up to 20-50% depending on the timing of divergence. We also find that these biases can affect the detection of interpopulation differences in recombination rate, causing both false positive and false negatives depending on the scenario. We discuss future possibilities for mitigating these biases and recommend that investigators exercise caution and confirm that their study populations meet assumptions before deploying these methods.

## Introduction

Recombination rate, the number of crossovers per unit genome per generation, plays a key role in shaping evolutionary processes and diversity in the genome. For example, through the action of linked selection, local rates of recombination are a chief determinant of patterns of genetic diversity throughout the genome (Begun & Aquadro, 1992; Burri, 2017; Cutter, 2019; Haddrill et al., 2014; Korunes et al., 2021). Genome-wide rates of recombination also modulate diverse processes such as adaptation, speciation, and introgression (Dapper & Payseur, 2017; Samuk et al., 2017; Schumer et al., 2018; Stapley et al., 2017). There is also a growing appreciation that recombination rate is itself a trait that varies and evolves (Dumont & Payseur, 2008; Hunter et al., 2016; Johnston et al., 2016; Ritz et al., 2017; Samuk et al., 2020; Stapley et al., 2017). Accordingly, there has been great interest in efficient and accurate methods for estimating recombination rates.

Current methods for estimating recombination rates fall into two broad classes of methods: direct and indirect (Peñalba & Wolf, 2020). Of the direct measures, the three most popular approaches are linkage mapping, gamete sequencing, and cytological methods. With classical linkage mapping, map distances between genetic markers are measured by quantifying recombinant markers in the context of a genetic cross or pedigree (Broman, 2010; Rastas, 2017). The resolution of this approach is limited only by marker density and the sample size of individuals, but larger sample sizes can be grueling to carry out in the laboratory or unavailable in some populations. Further, identifying suitable diagnostic mapping markers can be limiting in some cases (e.g. in a highly homozygous population, Broman, 2010). Direct sequencing of pools of recombinant gamete genomes from single individuals using long/linked read sequencing is a newer approach that alleviates many of the issues of traditional mapping, but still requires differentiated markers to score crossover events between homologous chromosomes (Dréau et al., 2019; Rommel Fuentes et al., 2019; Xu et al., 2020). Cytological methods bypass this requirement by directly visualizing recombination-associated protein complexes in cell populations undergoing meiosis (Peterson et al., 2019; Peterson & Payseur, 2021). However, cytological methods are limited by the spatial resolution at which such visualization can occur (e.g. the resolution of immunostained gamete karyotypes, Peterson et al., 2019).

Because all direct methods of measuring recombination rates are fairly laborious, there has been increased interest in indirect measures of recombination rate that leverage readily available population genetic data. Chief among these are model-based methods that infer rates of recombination from patterns of linkage disequilibrium (LD), (Auton & McVean, 2007; Chan, Song, et al., 2012; Kamm et al., 2016; Spence & Song, 2019). These methods attempt to estimate recombination rates by statistically fitting recombination rates (derived from population genetic models/simulations) to observed patterns of LD. Rather than inferring recombination rate directly, LD-based estimators infer a *population scaled recombination rate*, ρ = 4N_e_*r*, where N_e_ is the effective population size and *r* is the theoretical per-generation recombination rate. LD-based methods are attractive because they (1) generally only require population-scale genomic data and (2) are very fast, often only requiring several computational hours or less (Spence & Song, 2019) and (3) are informative of time-averaged population historical recombination rates (Gil McVean & Auton, 2007). Accordingly, LD-based estimates of recombination rates have become extremely popular, and now vastly outnumber direct measures in the literature (Peñalba & Wolf, 2020; Stapley et al., 2017). These methods have also begun to be used to perform interpopulation comparisons of recombination rates (Peñalba & Wolf, 2020; Stapley et al., 2017).

Like all models, LD-based estimators of recombination rate make a variety of simplifying assumptions about the populations of interest. For one, they generally assume that the populations/loci of interest are evolving largely neutrally and have reached population genetic equilibrium in a number of ways (Stumpf & McVean, 2003). In particular, most methods assume that the populations being studied have reached an equilibrium between recombination and population scaled mutation, such that LD accurately reflects patterns of recombination rate (G. McVean, 2007). Further, it is generally assumed that any form of selection that might distort patterns of LD (e.g. sweeps) has not recently occurred (Chan, Song, et al., 2012). Finally, these methods make the general assumption that demographic processes that distort genome-wide patterns of LD, such as population size changes, have not occurred (recall that ρ is directly dependent on N_e_, Auton & McVean, 2007).

While some of these assumptions may be robust to violation, work has shown that some violations can result in biased estimates. For example, (Dapper & Payseur, 2018) showed that recombination estimates from LDhat (Gil McVean & Auton, 2007) are highly sensitive to changes in population size. This can be ameliorated in some cases by incorporating known changes in population size into the estimation procedure, such as implemented in the software pyrho (Spence & Song, 2019).

Along with changes in population size and selection, another process that can greatly alter patterns of LD is gene flow. Gene flow and subsequent admixture between diverged populations can have complex effects on patterns of LD within each population (Nei & Li, 1973; Ohta, 1982). These effects range from large and genomically variable increases in LD due to segregation of divergent haplotypes, to genome-wide decreases in LD as populations become coupled and increase local N_e_ (Nei & Li, 1973; Ohta, 1982). While it is now widely accepted that gene flow is commonplace in natural populations (Barton, 2008; Mallet, 2005; Suvorov et al., 2021; Waples & Gaggiotti, 2006), and there has not been a systematic study of the effects of gene flow on LD-based measures of recombination. Further, it remains unclear how gene flow (or any other violation of assumptions) impacts our ability to detect differences in recombination rate *between* (as opposed to within) populations using LD-based methods.

Here, we address these issues using forward-time population genetic simulations. We attempt to answer two specific questions. First, how does gene flow between populations affect the precision and accuracy of LD-based estimates of recombination rate within populations? Secondly, how does gene flow affect our ability to detect evolved differences in recombination rate between populations? Our primary goal is to answer these questions in the context of a core set of realistic demographic scenarios, and not perform an exhaustive exploration of parameter space. Overall we hope to help investigators understand key sources of bias in LD-based estimates of recombination rate in natural populations and highlight areas of future development.

## Methods

### Code availability

All scripts used in the analyses described below are available as a repository on Github (http://github.com/ksamuk/LD_recomb).

### Forward time simulations with SLiM

To explore how the timing and amount of gene flow affect estimates of recombination rate, we performed forward-time population genetic simulations using SLiM version 3.3 (Haller & Messer, 2019). The basic form of all the simulations was an isolation-with-migration scenario: a single ancestral population diverges into two subpopulations with a static amount of bidirectional gene flow (Figure 1). Populations were composed of diploid individuals with 100kb genomes arranged in a single chromosome. We used genome-wide average estimates of effective population size, mutation rate, and empirical recombination rate from natural populations of *Drosophila melanogaster (Adrion, Cole, et al*., *2020)*: Mutation rate = 5.49×10^−9^ (Li & Stephan, 2006); Recombination rate = 2.23×10^−8^, (average of chromosome 2R, Comeron et al., 2012); N_e_ = 1.72M (Li & Stephan, 2006). Recombination and mutation rates were conservatively modeled as uniform across the 100kb genome. Following standard practice for forward-time simulations, all simulations were run with an *in silico* population size of N=1000, and simulated mutation and recombination rates scaled by a factor of N_e_/N as per the SLiM manual (Haller & Messer, 2019). Note that generation times are also subject to scaling, and for simplicity, we will refer to all generations in terms of back-transformed actual generations rather than SLiM generations (1 SLiM generation ≈ 1751 actual generations with our scaling factor).

**Figure 1.**
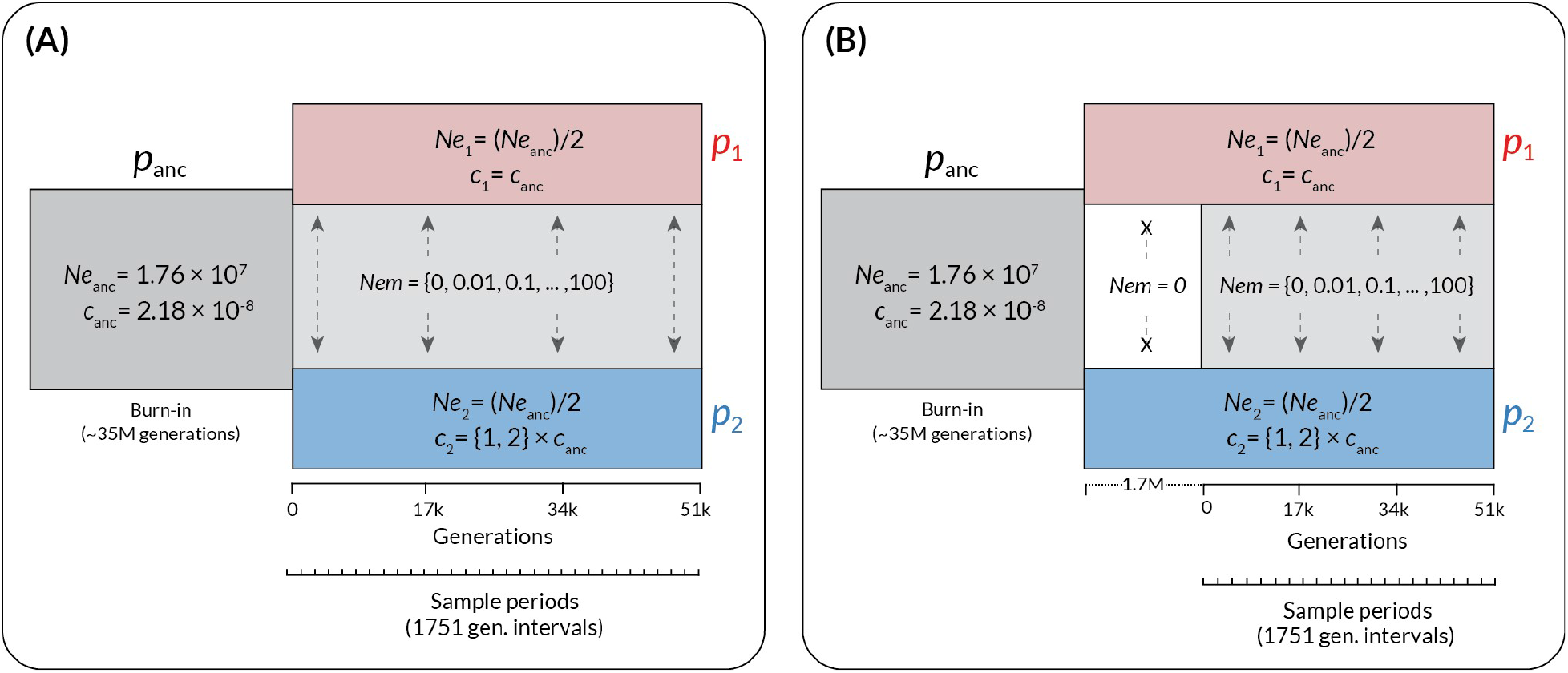
The structure of the forward-time simulations performed in SLiM. Time in back-transformed generations is shown along the x-axis, and the populations in existence at a given time are shown as rectangles. *p*_anc_ = the ancestral population, *p*_1_ = the subpopulation with unchanged recombination rate, and *p*_2_ = the subpopulation with increased recombination rate (if applicable). Effective population sizes (N_e_) and recombination rates (*c*, in units of cM/Mb) are shown for each population, with the values for the subpopulations shown relative to the ancestral value. Variable elements of the simulation are shown in braces. Time in generations post-divergence is indicated below the plots, with the pre-contact isolation period in (B) shown as a dotted line preceding the main axis. Sample periods indicate intervals at which genotypes were output for analysis.

### Parameter space

To explore how variation in gene flow affects estimates of recombination, we varied the amount of gene flow over five orders of magnitude: 0, 0.01, 0.1, 1, 10, 100, in standard units of N_e_m (the product of the effective population size and the migration rate). These values were chosen to encompass total isolation (N_e_m = 0), limited gene flow (N_e_m = 0.01-0.1), moderate gene flow in interconnected metapopulations (N_e_m = 1-10, Morjan & Rieseberg, 2004; Waples & Gaggiotti, 2006), and a scenario of a nascent hybrid swarm (N_e_m = 100). We also varied the timing of the onset of gene flow, with gene flow beginning either immediately after divergence or after a period of isolation. We performed preliminary simulations to determine a period of isolation (∼1.7M generations in our case) that produced levels of genomic divergence (Figure S1) similar to those observed in natural population pairs that exhibit genome-wide genetic divergence but still actively exchange genes (F_ST_ ∼ 0.4, Morjan & Rieseberg, 2004; Roux et al., 2016). Finally, to explore how gene flow impacts the detection of population *differences* in recombination rate, we modeled scenarios where recombination rate either remains constant in both subpopulations or instantaneously increases by a factor of 2 at the time of divergence in one of the two subpopulations (always subpopulation two). This magnitude of this difference is well within the range of variation in recombination rate reported for a wide variety of species (Stapley et al., 2017). In biological terms, an instantaneous increase in population recombination rate could be readily mediated by an environmental change (e.g. temperature, Lloyd et al., 2018) or via a change in mating system (Brandvain & Wright, 2016). We note that this instantaneous change is a “best case” scenario for detecting interpopulation differences in recombination rate, and thus any loss of power to detect differences in recombination that occurs due to gene flow will be conservative.

### Details of demographic events

Each simulation began with a single population of size Ne_anc_, which evolved for a 35M generation burn-in period (following the general practice of a 10-20 N_e_ burn-in period, Haller & Messer, 2019). This initial period was followed by divergence into two subpopulations, each with size Ne_anc_/2. Gene flow (for cases where N_e_m > 0) began immediately at the time of divergence or after a 1.7M generation period of isolation and was symmetrical in magnitude and bidirectional. Changes in recombination rate occurred at the time of divergence and instantaneously applied to all individuals in subpopulation two only.

Starting at the time of divergence and thereafter in intervals of 1751 generations, we collected a random sample of 25 individuals (a total of 50 haploid genomes) from each population and saved their complete genotypic at all sites in VCF format. We stopped the simulations after 51 000 generations. Each parameter combination was replicated 100 times, for a total number of ∼n=48 000 population samples.

### Estimation of recombination rate using pyrho

While there are a variety of LD-based estimators of recombination rate, we elected to use pyrho (Spence & Song, 2019) for estimation in this study. It shares its statistical foundation with the most widely used LD-based estimators (LD-hat & LD-helmet; Chan, Jenkins, et al., 2012; Gil McVean & Auton, 2007) while also having the ability to account for changes in effective population size such as we are modeling here (Spence & Song, 2019). As such, any estimation biases caused by gene flow will likely affect those approaches as least as much they affect pyrho. Direct comparisons with other methods are complicated by the fact that pyrho is the only model-based method that adequately accounts for changes in effective population (ReLERNN being a possible exception Adrion, Galloway, et al., 2020)

We followed the recommended practices for inferring recombination rate using pyrho (https://github.com/popgenmethods/pyrho). We parameterized the initial lookup tables using the effective population size and mutation rates used in the simulations (*unscaled* in this case). To account for changes in effective population size, we created lookup tables that accounted for a change of N_e_/2 (1.72M to 8.6M) in time steps of 1751 generations in the past. This allowed us to have an appropriately timed lookup table for each step of the simulation. We used the built-in methods to infer the hyperparameters of window size (best fit 100) and block penalty (best fit 1000). Using this baseline, we inferred recombination rates using the genotype data (VCF format) from both subpopulations at each time point, for a total of ∼96 000 pyrho fits. All computation was performed using the Duke University Computing Cluster, running CentOS Version 8.

### Statistical analyses

We performed all data processing and visualization using the tools of the tidyverse package in R 4.0.3 (R Core Team, 2018; Wickham, 2017). To examine how gene flow between populations affects the accuracy of LD-based estimates of recombination, and the context of the various factors explored in our simulations, we performed an analysis of variance using a linear mixed model with Gaussian errors fitted via the lmer() function from the lme4 package (Bates et al., 2007). This model had the following form: Recombination rate = (1|simulation replicate) + (1|simulation generation) + gene flow magnitude + recombination rate change, where (1|[factor]) denotes a random intercept and “:” denotes an interaction. All variables were standardized (mean-centered and scaled by standard deviation) prior to analysis. To simplify interpretation, we fitted separate models for the continuous gene flow and secondary contact scenarios.

## Results

### Inference when recombination rate is identical between populations

When the recombination rate remained constant between diverging populations, we found that gene flow introduced two types of systematic biases in estimates of recombination rate within populations (Figure 2A). These effects began when N_e_m >= 1 in both the continuous gene flow and secondary contact models. First, in the model of continuous gene flow, when N_e_m >= 1, we observed a systematic increase (overestimate) in estimated rates of recombination in both populations (Figure 2A, 2B, top row, N_e_m = 1-100). This increase was statistically significant (Type III Wald chi-square = 5090.07, p < 2.0×10^−16^; coefficient for gene flow = 0.63–0.67 (95% CI), t(19495) = 71.34, p < 0.001). When the migration rate was moderate to high (N_e_m 10-100), the recombination rate was overestimated by approximately 10-20% (Figure 2B). This effect is consistent with migration causing the populations to become coupled, behaving as a single population with a larger Ne and thus inflating the population-scaled estimate of recombination rate.

**Figure 2.**
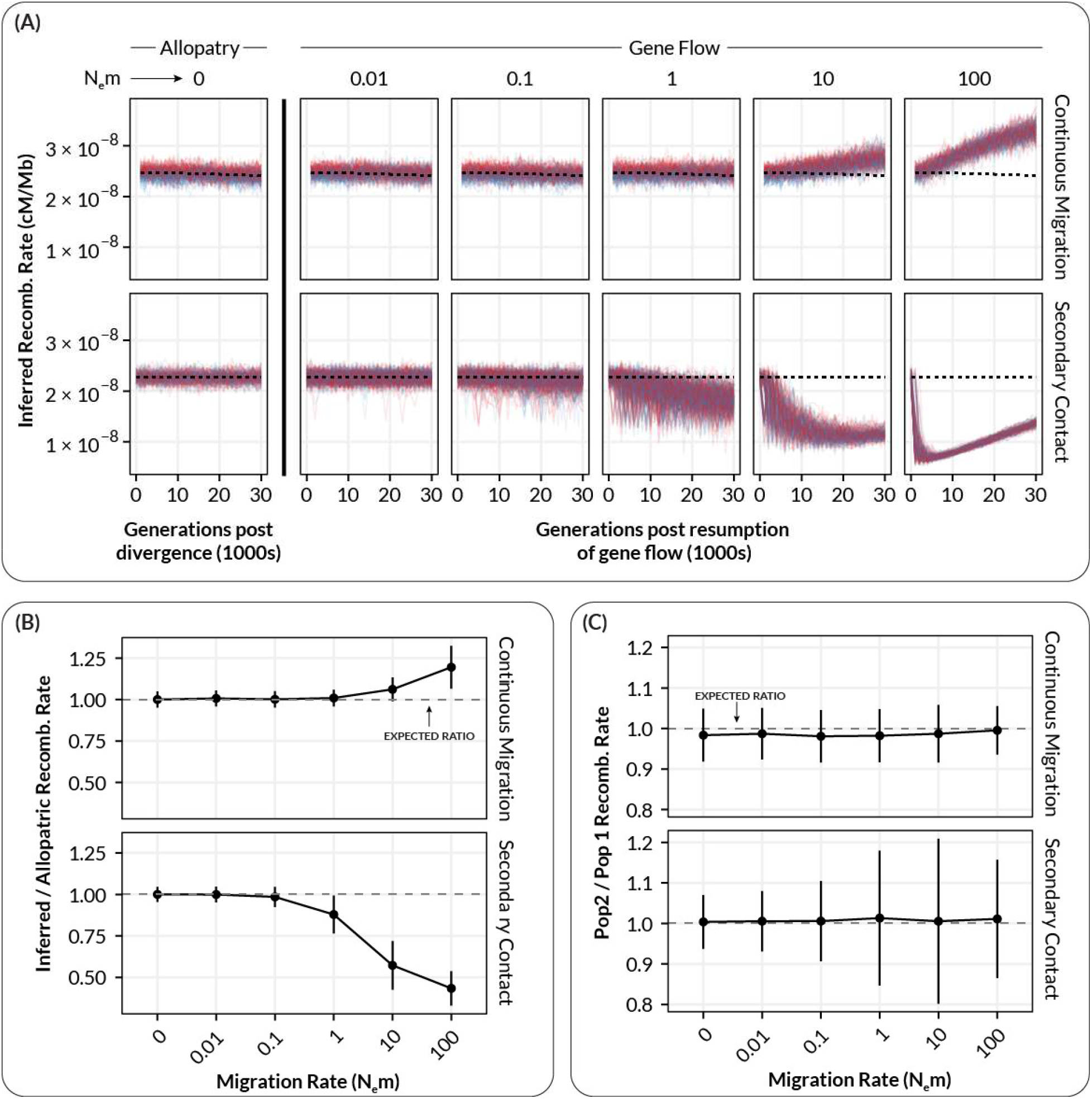
The relationship between inferred recombination rate and the migration rate in simulated populations where recombination rate remains constant in both subpopulations. (A) Inferred recombination rates for individual simulations at varying levels of migration. Each plot shows inferred rates for simulation replicates (transparent lines) of population 1 (red, unchanged recombination) and population 2 (blue, increased recombination) for a single migration rate. Dashed lines show the expected inferred value in the absence of gene flow (inferred from N_e_m = 0). (B) Summarized inferred recombination rates (y-axis) for each level of migration (x-axis) from the simulations in A. Points are mean values and error bars depict standard deviations (summarized across all generations). Dashed lines show the expected inferred value in the absence of gene flow for each population (i.e. the mean value for N_e_m = 0). (C) The inferred *difference* in recombination rate between population 1 and population 2 (*p*_*2*_ -*p*_*1*_) as a function of migration rate. Points and errors bars are as in B.

In contrast to the continuous gene flow case, under a model of secondary contact, there was a marked systematic decrease (underestimate) of recombination rates, which also became visible when N_e_m >=1 (Figure 2A, 2B, bottom row, N_e_m = 1-100). This decrease was statistically significant (Type III Wald chi-square = 1512, p < 2.2×10_-16_; coefficient for gene flow = -(0.54–0.49) (95% CI), t(31846) = -38.88, p < 0.001). The magnitude of this decrease was substantial: on average, populations experiencing N_e_m = 1 had recombination rates about 20% lower than expected, with this increasing to 50% when N_e_m = 10 or higher (Figure 2A, Figure 2B). This decrease was accompanied by a statistically significant increase in the *variance* of recombination rate estimates, especially for N_e_m = 1-10 compared to N_e_m < 1 (Fig 2A, bottom row; F-test for equivalency of variance, F(10429,13860) = 0.20863, p< 2.2×10^−16^). A systematic increase in the mean and variance of LD within populations is consistent with allele frequency differences between populations manifesting as migration-associated LD, and deflating estimates of recombination rate. When gene flow was very high, there was a visible recovery of estimated recombination rates (Figure 2A, bottom row, N_e_m = 100), presumably due to migration homogenizing allele frequencies and increased effective population sizes increasing the rate at which recombination breaks down migration-associated LD.

When comparing recombination rates between *p*_*1*_ and *p*_*2*_, the “coupling” bias observed in the continuous migration scenario did not appear to systematically affect the *ratio* of recombination rate between the two populations (Figure 2C, Continuous Migration).

However, in keeping with the previous result, migration-associated LD in the secondary contact model appeared to greatly increase the variance in the ratio of recombination rates between populations when N_e_m >= 1 (Figure 2C, Secondary Contact).

### Inference when recombination rate differs between populations

When recombination rates diverged between populations, we also observed the two forms of bias described above (Figure 3). The estimates from the continuous gene flow scenario exhibited a statistically significant increase (Type III Wald chi-square = 8936.44, p < 2.2×10^−16^; coefficient for gene flow =0.65-0.67 (95% CI), t(19495) = 94.53, p < 0.001) whereas estimates from the secondary contact model exhibited a statistically significant decrease (Type III Wald chi-square = 1512, p < 2.0×10^−16^; coefficient for gene flow = -(0.27-0.22) (95% CI), t(34505) = -23.22, p < 0.001). However, the results differed from simulations with constant recombination rates in a number of important ways. First, there was a clear difference between the continuous migration and secondary contact models in the overall trajectory in the population-specific estimates of recombination rate (Figure 3A). In the continuous gene flow models, there was an overall positive trend for the estimates of recombination rate in *p*_*2*_ even in the absence of gene flow (Figure 3A, continuous migration). This was presumably caused by a lag in the establishment of equilibrium levels of LD within *p*_*2*_ that reflect the new recombination rate (which spontaneously changed at the time of divergence). This lag resulted in the recombination rate in *p*_*2*_ being consistently underestimated (because it had not reached its new equilibrium), in addition to the coupling effect observed previously (Figure 3B and 3C, continuous migration).In the case of the secondary contact model, we did not observe the same positive trend for recombination rate estimates in *p*_*2*_, likely because the isolation period (1.7M generations) was sufficiently long enough for *p*_*2*_ to establish an equilibrium level of LD prior to secondary contact (Figure 3A, Secondary Contact).

**Figure 3.**
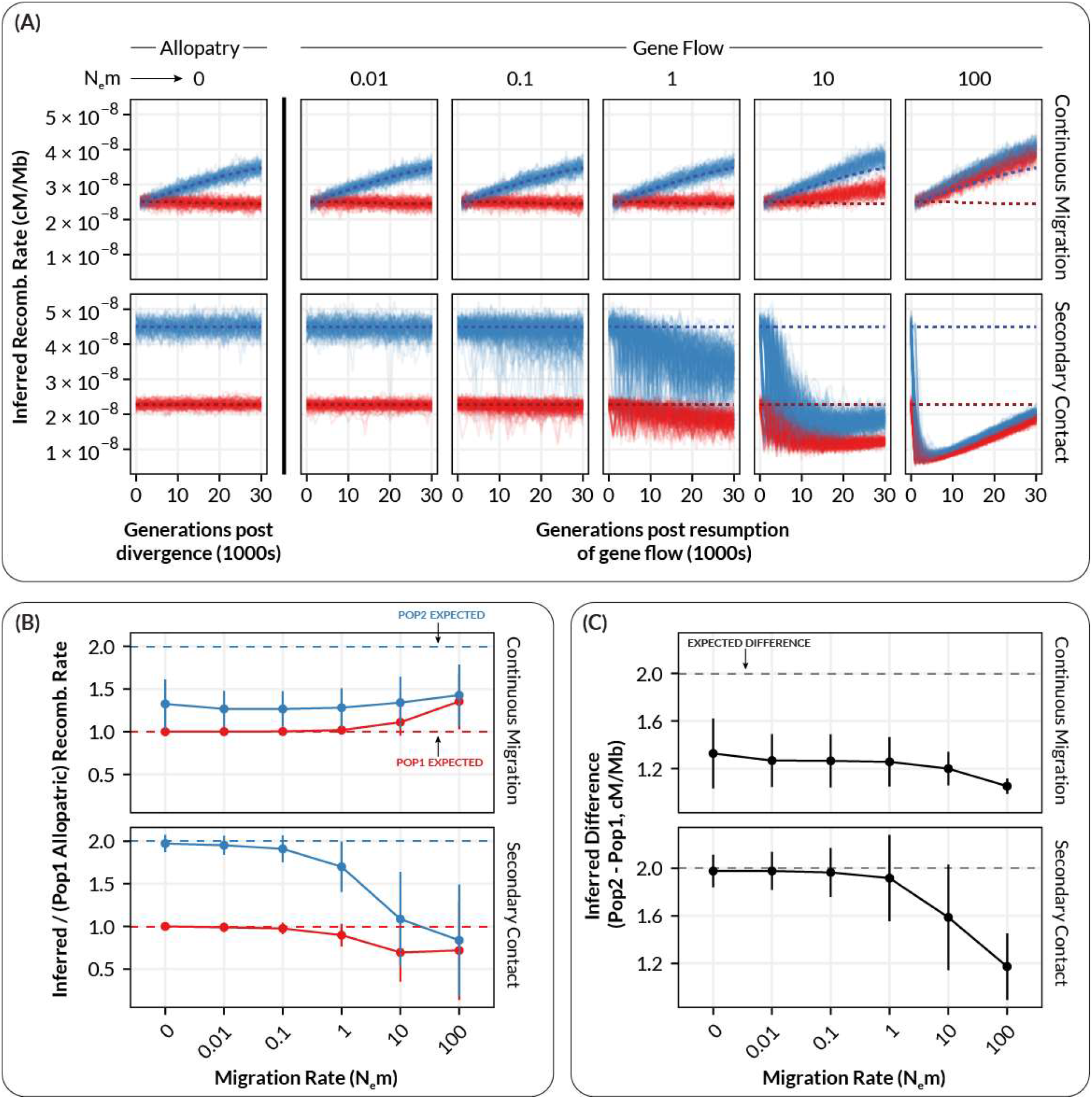
The relationship between inferred recombination rate and the migration rate in simulated populations where recombination rate increases by a factor of two in one subpopulation. (A) Inferred recombination rates for individual simulations at varying levels of migration. Each plot shows inferred rates for simulation replicates (transparent lines) of population 1 (red, unchanged recombination) and population 2 (blue, increased recombination) for a single migration rate. Dashed lines show the expected inferred value in the absence of gene flow (inferred from N_e_m = 0). (B) Summarized inferred recombination rates (y-axis) for each level of migration (x-axis) from the simulations in A. Points are mean values and error bars depict standard deviations (summarized across all generations). Dashed lines show the expected inferred value in the absence of gene flow for each population (i.e. the mean value for N_e_m = 0). (C) The inferred *difference* in recombination rate between population 1 and population 2 (*p2* -*p*_*1*_) as a function of migration rate. Points and errors bars are as in B.

In keeping with the scenario with constant recombination rates, starting at N_e_m ∼ 1, migration-associated LD resulted in the systematic underestimation and increase in variance for estimated recombination rates within both *p*_*1*_ and *p*_*2*_ (Figure 3B, Secondary Contact; Type III Wald chi-square = 538.97, p < 2.0×10^−16^; coefficient for gene flow = - (0.27-22) (95% CI), t(34505) = -23.22, p < 0.001; F-test for equivalency of variance, F(7734,13579) = 0.2174, p< 2.2×10^−16^). In addition, the observed divergence in recombination rate between *p*_*2*_ and *p*_*1*_ (which was always expected to be +2 cM/Mb) decreased with increasing levels of gene flow (Figures 3B and 3C, Secondary Contact). This effect would likely result in an increase in false negatives with increasing gene flow (i.e. finding no difference in recombination rate between populations when there is in fact one). This decrease in the observed divergence between populations is again likely the outcome of the population-specific levels of LD becoming coupled/merged at moderate to high levels of gene flow, resulting in the populations exhibiting LD (and hence recombination rate estimates) intermediate to what would be expected in the absence of gene flow.

## Discussion

Accurate estimates of recombination rate are key to understanding the causes and consequences of recombination rate variation in natural populations. With the increasing availability of genome-wide sequencing data, LD-based estimators of recombination rate have become widely used in a large variety of taxa. However, while gene flow is widely known to shape patterns of LD in populations, the effect of gene flow on LD-based estimators of recombination rate remains largely unexplored. Here, use forward-time simulations to show that (1) gene flow can introduce substantial bias into LD-based estimates of genome-wide recombination rate and (2) the nature of this bias depends on the demographic and evolutionary history of the populations in question.

Our results here are consistent with theoretical predictions that gene flow between populations can affect LD: increasing in the magnitude and variance of LD at low migration rates as well as reducing LD via the “coupling effect” we observed at higher rates of gene flow. Our study shows how these predictions play out with modern methods and genomic data, and also provides a sense of the magnitude of the potential degree of misestimation - in our case, ranging from 20-50 percentage points in cases of moderate gene flow. For comparison, a recent study of population-level differences in recombination rate in *Drosophila pseudoobscura* revealed genetically based interpopulation differences on the magnitude of approximately 10% measured using replicated linkage maps in each population (Samuk et al., 2020). Using LD-based estimators, an observed difference of this magnitude could be spuriously generated by modest levels of gene flow alone, or missed altogether due to coupling at higher level gene flow is high. In addition, the specific magnitude and direction of the bias introduced by gene flow is difficult to know without precise knowledge of the population/demographic histories of the populations in question. This should give pause to anyone planning on using LD-based methods to infer recombination rate in non-equilibrium populations.

One key question is whether there are methods to control for or counteract the increased variance and/or biases in the estimation of recombination rate caused by gene flow. One approach could be to identify and remove introgressed haplotypes from datasets prior to inferring recombination rate, thereby removing migration-associated LD. This would require “pure” samples from the source populations, such that the population of origin could be assigned to haplotype blocks (Dias-Alves et al., 2018). However, this method would only work if gene flow is infrequent enough that coupling (of both LD and allele frequencies) has not occurred. The upward bias and increased variance in recombination rate that occurs as a result of coupling, together with the homogenization of allelic differences between populations at higher levels of gene flow will likely make a “filtering” scheme very difficult (perhaps impossible) to achieve. One approach may be to attempt to jointly estimate a demographic model along with population-specific recombination rates, as has been done with mutation rates (DeWitt et al., 2021). However, given the existing complexity and uncertainty in inferring demographic models, we suspect it may be difficult to disentangle the complex interdependencies between gene flow, population size, and estimates of recombination rate.

Together with previous work (Dapper & Payseur, 2018), our results suggest that LD-based estimates of recombination rate need to be interpreted with great caution when studying non-equilibrium populations. Indeed, these methods are likely only appropriate when populations can be assumed to be evolving in the absence of any gene flow, and have reached a reasonable demographic equilibrium. However, it is now widely appreciated that gene flow is ubiquitous in natural populations (Ellstrand & Rieseberg, 2016; Waples & Gaggiotti, 2006). This may mean that many published LD-based estimates of recombination rate are incorrect. Without empirical maps to compare existing LD-based estimates, it is difficult to say just how incorrect. What can be said is that the levels of gene flow required to introduce non-trivial biases into estimates of recombination rate, i.e. N_e_m ∼1-10, are not uncommon in natural populations (Slarkin, 1985; Waples & Gaggiotti, 2006). It is also worth noting that it is not the case that two populations being studied have to be exchanging genes themselves (e.g. which would not the case when studying two reproductively different species), but just that one or more of the populations are exchanging genes with some *other* population (e.g. an unsampled population of the same species).

If many LD-based estimates are incorrect, why do published LD-based estimates of recombination rate correlate well with direct estimates, e.g. from genetic maps? (Chan, Jenkins, et al., 2012; Gil McVean & Auton, 2007; Smukowski Heil et al., 2015). There are several considerations. First, the correlations that have been reported are by no means perfect (e.g. ∼Spearman’s Rho of 0.6: Smukowski-Heil et al. 2015; r^2^ = 0.37-63: Chan, Song, et al., 2012) and depend greatly on the genomic scale at which they are measured (Smukowski-Heil et al. 2015). Second, simple correlations between LD-based and empirical estimates cannot detect genome-wide differences in the estimates of recombination rate, such as those due to the coupling effects we observed. Such effects would be visible as differences in the *intercept* of a linear regression, rather than the R^2^, for example. Finally, the species where these correlations have been examined (humans and *Drosophila melanogaster*) may meet the assumptions of demographic equilibrium more readily (Ochoa & Storey, 2019; Suvorov et al., 2021). While such assumptions may be reasonable for these populations, for which LD-based estimators were originally developed, they are much less likely to hold in many natural populations. Notably, they are likely rarely met in populations that have recently adaptively diverged in the presence of gene flow, which have lately been the subject of increased research interest (Linck & Battey, 2019; Ravinet et al., 2017). The equilibrium assumption is likely not valid in populations in which the recombination rate has recently changed (Brandvain & Wright, 2016), reducing the utility of these estimates for studying the rapid evolution of recombination rates.

While we only focused on a single implementation of one type of LD-based estimator of recombination (pyrho), it is likely that other population genetic methods will also suffer from the effects we describe here. LD is the “information” used by all estimators, either directly as in methods like LDjump (Hermann et al., 2019) or indirectly as in machine learning methods like ReLERNN (Adrion, Galloway, et al., 2020). That said, in the case of the latter method, it may be possible to overcome some of the issues we’ve identified if the training datasets were simulated with an accurate demographic model. As such, the distorting effects of gene flow on LD need to be carefully considered when applying any statistical methods for inferring recombination rate approaches. We also stress that our simulations do not suggest that LD-based estimators and their implementations are wrong per se, but rather that the assumptions under which LD-based estimates are biologically accurate are readily violated by levels of gene flow and divergence common seen in natural populations.

## Conclusion

Studying variation in recombination rate is difficult. LD-based methods for inferring recombination rate are attractive in their data requirements, but require strong assumptions to be met. As we have shown here, gene flow readily violates these assumptions and introduces biases and decreases in precision, in a variety of ways that are difficult to identify in a given study population. This is problematic because gene flow is extremely common in natural populations. How should we proceed? Rather than attempt to squeeze blood from the proverbial stone, we believe that the most straightforward solution to the problems we outline here is simply to prioritize the use of direct, empirical methods for measuring of recombination rate. This decision is made hopefully simpler with the increased ease and low cost of creating traditional linkage maps and performing gamete sequencing. That said, LD-based approaches remain important tools for hypothesis generation, and when paired with direct estimates of recombination rate can provide a detailed picture of both the past and present landscape of recombination rates in natural populations.

## Acknowledgements

Support this project was provided by National Science Foundation grants DEB-1545627, 1754022, and 1754439 to MAFN. KS was additionally supported by a Natural Sciences and Engineering Research Council of Canada Postdoctoral Fellowship. We thank members of the Noor lab and Dr. Katharine Korunes for helpful discussions and for providing comments on an early draft of this paper.

## Supplemental Material

**Figure S1.**
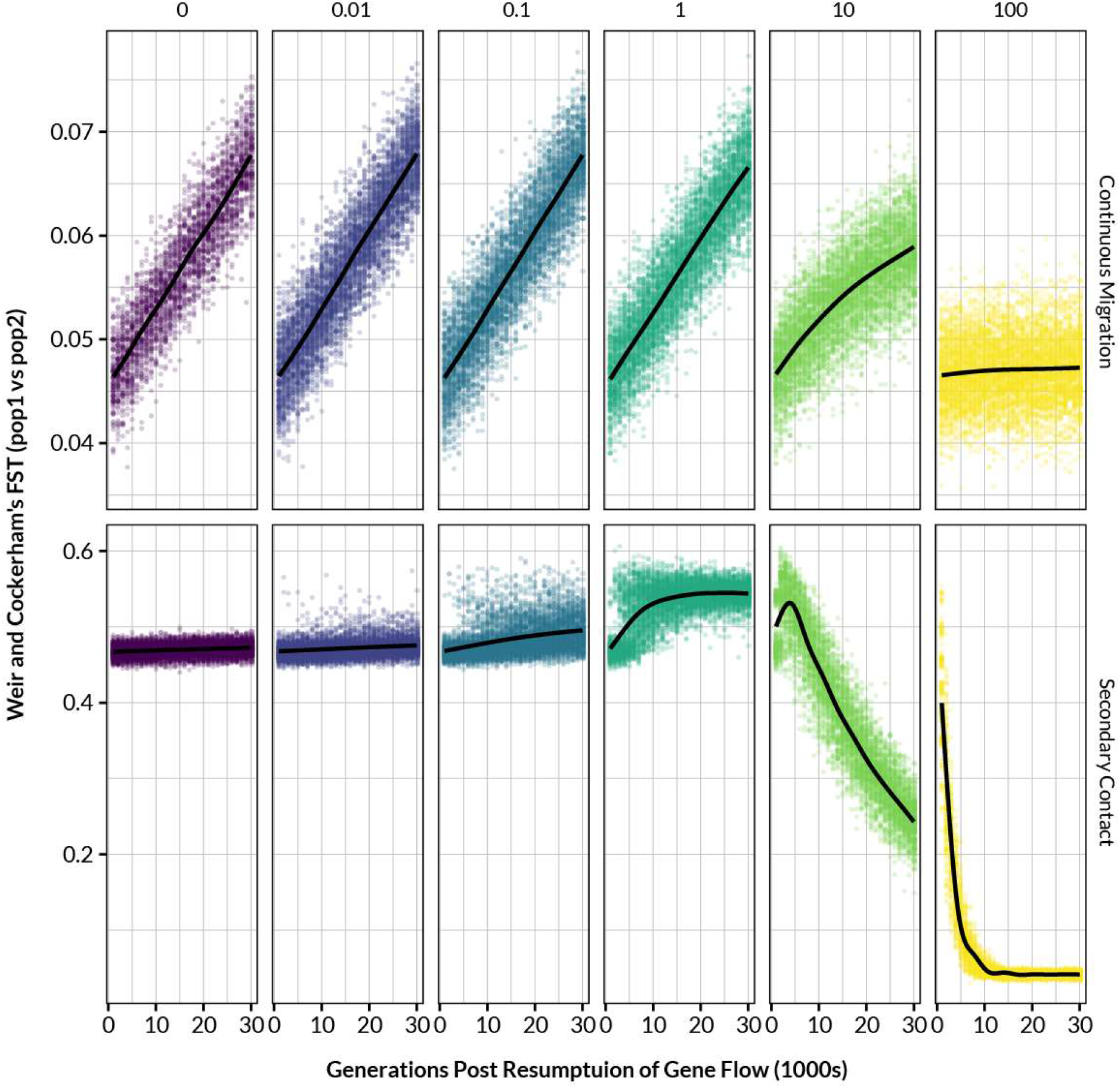
Weir and Cockerham’s FST between simulated populations as a function of time in generations under various combinations of migration rate (columns, N_e_m) and isolation scenario (rows). Black lines are smoothed LOESS fits. Note the difference in y-axis scales between the rows.

